# Precise triggering and chemical control of single-virus fusion within endosomes

**DOI:** 10.1101/2020.04.12.038109

**Authors:** Sourav Haldar, Kenta Okamoto, Peter M. Kasson

## Abstract

Many enveloped viruses infect cells within endocytic compartments. The drop in pH that accompanies endosomal maturation, often in conjunction with proteolytic factors, serves as a trigger for viral fusion proteins to insert into the endosomal membrane and drive fusion. The dynamics of this process has been studied by tracking viruses within living cells, which limits the precision with which fusion can be synchronized and controlled, and by reconstituting viral fusion to synthetic membranes, which introduces non-physiological membrane curvature and composition. To overcome these limitations, we have engineered the chemically controllable triggering of single-virus fusion within endosomes. We isolate influenza virus:endosome conjugates from cells prior to fusion, immobilize them in a microfluidic flow cell, and then rapidly and controllably trigger fusion. This platform demonstrates lipid-mixing kinetics that are grossly similar to influenza fusion with model membranes but display some subtle differences. Because it preserves endosomal membrane asymmetry and protein composition, it also provides a means to test how perturbations to endosomal trafficking and cellular restriction factors affect viral membrane fusion.

## Introduction

Many enveloped viruses infect host cells via a process of membrane fusion between the viral envelope and endosomal membranes [1,2]. Influenza is perhaps the prototypical enveloped virus, where entry occurs within mid-late endosomes and is triggered by endosomal acidification during the trafficking process. Acidification drives a series of change in the hemagglutinin coat protein that lead to exposure and insertion of a fusion peptide in the endosomal membrane, refolding of the hemagglutinin ectodomain, and membrane fusion [3–5]. Opening of a fusion pore makes the viral interior topologically continuous with the host cell cytoplasm, and subsequent viral uncoating [6] permits release of viral RNA and replication (Fig. 1(A)).

**Figure 1.**
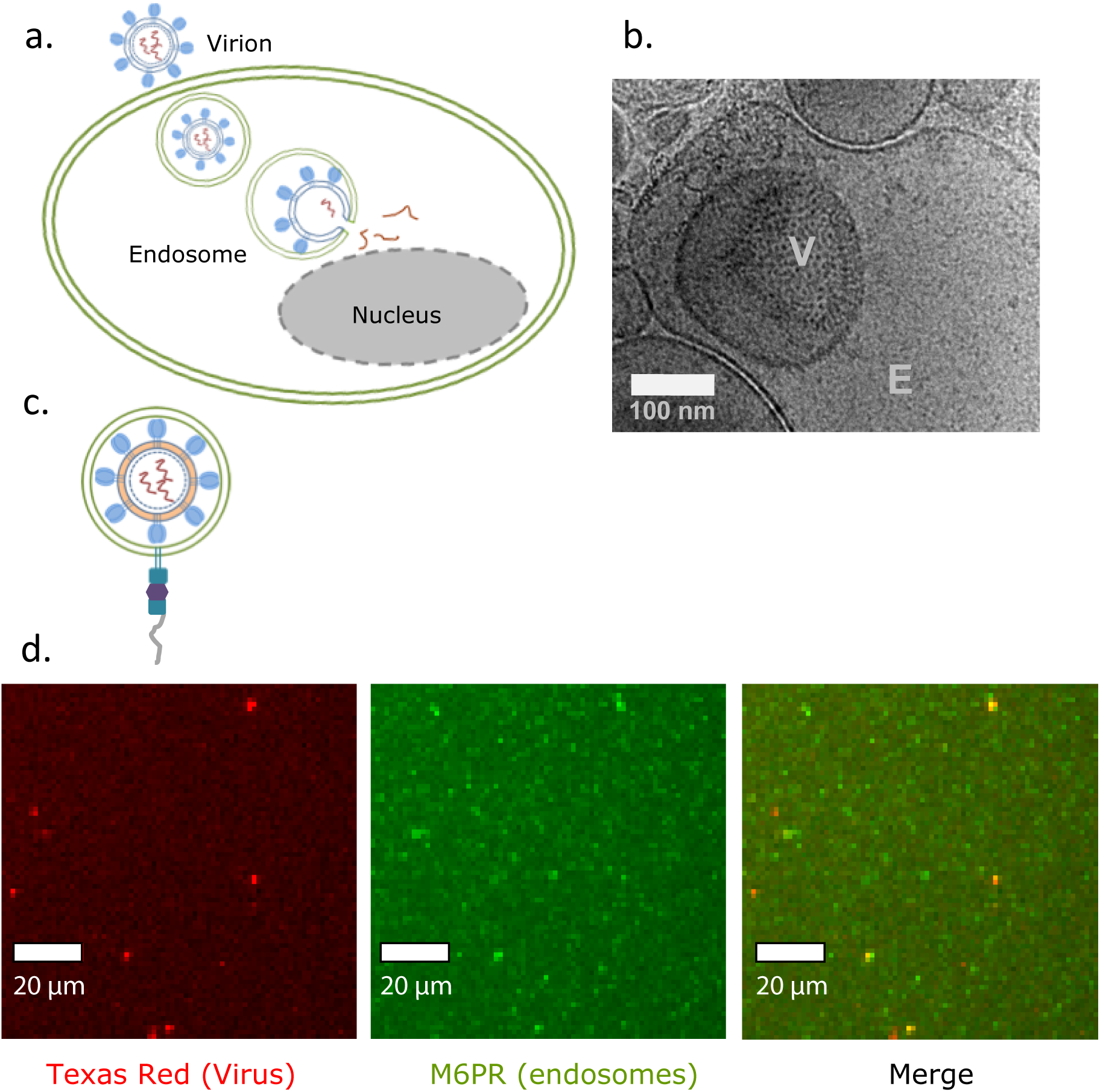
Isolation of virus containing endosomes. (a) Schematic representation of influenza virus entry into the host cell. Upon binding to the plasma membrane of the host cell, influenza is internalized by endocytosis and contained in early endosomes. Endosomal maturation is accompanied by luminal acidification. This drop in pH triggers fusion between viral and endosomal membranes, driven by influenza hemagglutinin, which in turn permits release of viral RNA into the host cytoplasm. (b) Representative cryo-EM image of a likely influenza virus (marked V) enclosed in an endosome (marked E). (c) Schematic representation of isolated, surface-immobilized virus:endosome conjugates prior to triggering of fusion. The viral membrane is fluorescently labeled with Texas Red (TR) at self-quenching concentration to permit subsequent monitoring of fusion. The virus:endosome conjugate is immobilized by incorporation of biotin-PE and then binding to avidin:PLL-PEG-Biotin on a passivated glass coverslip. See Methods for details. (d) Representative epifluorescence microscopy images of virus:endosome conjugates. The red channel shows Texas Red in the viral membrane, and the green channel shows M6PR stained with a rabbit primary antibody and an Alexa488-conjugated secondary antibody. The merged image shows co-localization of virions (red) and endosomes (green) as yellow particles.

While initial insights into viral trafficking and fusion were obtained via bulk experiments [7–12], measurement of fusion by individual virions has permitted more precise characterization of the pathways and mechanisms by which viral entry occurs. Such single-virus fusion measurements have been made observationally in living cells or in a chemically controllable manner between viruses and engineered target membranes [13–21]. To better examine the effect of endosomal composition and geometry, we report a new strategy to measure single-virus fusion within endosomes in a controllable manner. To achieve this, we isolated influenza-containing endosomes from cells prior to fusion, immobilized them in a microfluidic flow cell, and monitored lipid mixing upon precise retriggering of fusion. This single-virus:endosome fusion assay permits direct comparison between viral fusion within endosomes and fusion to the exterior of liposomes or planar bilayers, where the membrane geometry and composition are not maintained. It thus helps elucidate the mechanistic determinants of fusion within cells as well as the role of cellular factors on the influenza virus:endosome fusion. The approach is easily generalizable to any enveloped virus that undergoes pH-triggered entry (such as dengue, Zika) and likely to other viruses such as Ebola and SARS that require pH in conjunction with endosomal factors to enter.

### Chemicals, Buffers and solutions

Phosphate-buffered saline: 10 mM NaH_2_PO4, 150 mM NaCl. The following buffers were used while inhibiting endosomal acidification: PBSN: phosphate-buffered saline, 30 mM ammonium chloride. CBN: 3mM imidazole, 1 mM EDTA, 30 mM ammonium chloride, pH 7.4. Homogenization buffer (HBN): 3mM imidazole, 1 mM EDTA, 30 mM ammonium chloride, 250 mM sucrose, HBN (+): 3mM imidazole, 1 mM EDTA, 30 mM ammonium chloride, 250 mM sucrose, protease inhibitor cocktail and 30 µM cyclohexamide. All sucrose solutions were made in CBN buffer, pH 7.4 and contained protease inhibitor cocktail and 30 µM cyclohexamide (mixed just before use). Fusion Buffer: phosphate-buffered saline, 90 mM sodium citrate, 10 µM FCCP, pH 5.0. All sucrose solutions were incubated with charcoal (10% w/v) for two hours with constant stirring to remove any impurities and subsequently pH-checked.

### Cell culture

Baby hamster kidney cells, (BHK Strain 21) were cultured in Dulbecco's Modified Eagle Medium (DMEM) supplemented with 10% heat-inactivated fetal bovine serum FBS (BioWest), 10 mM HEPES, and 1% Penicillin-Streptomycin solution (10,000 units penicillin and 10 mg streptomycin/mL) in a humidified incubator at 37 °C, with 5% CO_2_.

### Influenza virus

Influenza A virus (strain X-31, A/Aichi/68, H3N2) was purchased from Charles River Laboratories (Wilmington, MA). For lipid mixing experiments, virus was labeled with the lipophilic dye Texas Red-DHPE at a self-quenched concentration as described previously [18].

### Isolation of endosomes from BHK-21 cells

Endosomes containing viral particles were isolated from BHK cells in the following manner. After blocking acidification by incubating in 30 mM ammonium chloride in DMEM for 1 hour, a confluent monolayer of cells was infected with Texas Red (TR)-DHPE labelled X-31 virus. Binding of viral particles to the cells were carried out at 4 °C. Following which the cells were incubated at 37 °C for 20 min to allow endocytosis of viral particles. Cells were then washed two times with PBSN buffer to remove unbound viral particles. Next, the cells were scraped and sedimented in PBSN buffer by centrifugation at 4°C, 600 g for 5min. The pellet was resuspended in cold PBSN and re-pelleted by centrifuging at 4°C, 600 g for 5min. This step was repeated once with HBN and once with HBN (+). Finally, the pellet was resuspended in 3ml of cold HBN (+). Pelleted cells were homogenized and centrifuged to separate nuclei from post nuclear supernatant (PNS). The cell pellet was homogenized using a ball bearing homogenizer (28 µm clearance, Isotech, Heidelberg, Germany) on ice. The pellet was extruded at least 50 times. Homogenization was confirmed by making sure that no intact cells were visible upon microscopic examination of an aliquot. The homogenate was centrifuged at 4 °C, 2000g for 10min. The post nuclear supernatant (PNS), which contains cell organelles, was collected and maintained at 4°C. Endosomes were enriched from PNS by discontinuous sucrose gradient centrifugation: 2.5 mL PNS was mixed with 3 mL of 62% (w/w) sucrose in a 14 mL ultracentrifuge tube (Beckman). Next, 3ml of 35% and then 2 ml 25% sucrose solutions were layered above. The tube was with by 8% sucrose in HBN (+). The gradient was then centrifuged at 4 °C, 210,000 g for 1.5 hours. The endosomal fraction was identifiable as a milky white band at the interface between 35% and 25% sucrose as reported previously [22,23]. We chose mannose 6-phosphate receptor (M6PR) as a marker for mid-endosomes where influenza fuses, as reported previously [24]. 1 mL fractions were collected from top of the tube and stored at 4 °C until the next step. Fractions containing viral particles and endosomes were confirmed by immunoblotting (Fig. S1). Relevant fractions were further concentrated by centrifuging at 4 °C, 100,000 g for one hour in CBN. Pelleted endosomes were resuspended in 80 µl CBN buffer and kept at 4 °C until surface-immobilization and fusion.

### Single virus lipid mixing assay

All measurements were carried out inside a microfluidic flow cell as described previously [25]. Briefly, a glass coverslip inside a PDMS flow cell was coated with 95% PLL-PEG: 5% PLL-PEG-biotin. The biotin layer was functionalized by incubating with a solution of avidin. Endosomes were biotinylated by addition of 0.005% DPPE-biotin in solution and captured inside the flow cells by incubating 5 µl of endosome suspension for 30 min at 4 °C. Biotin-avidin binding enabled tethering of endosomes to the glass surface of the flow-cell. Any unbound or loosely bound particles were removed by extensive rinsing of the flow cell with CBN. Low pH buffer was infused, and single virus-endosome lipid mixing events at 37 °C were monitored by fluorescence microscopy. Virus-liposome measurements were carried out analogously using 100-nm extruded liposomes composed of 67% POPC: 20% POPE: 10% cholesterol: 2% GD1a: 1% DPPE-biotin as described previously [25].

### Microscopy

All epifluorescence micrographs and videos were acquired with a Zeiss AxioObserver using a 20 X air objective, NA= 1.49 (Zeiss), equipped with a Spectra-X LED Light Engine (Lumencor, Beaverton, OR) as an excitation light source, and additional excitation/ emission filter wheels. Images were recorded with an Andor Zyla sCMOS camera (Andor Technologies, Belfast, UK) using 16-bit image settings, and were captured with Micromanager software. Lipid mixing kinetics of virus:liposome and virus:endosome conjugates from video microscopy were analyzed via MATLAB scripts (The MathWorks, Natick, MA) previously reported [18] (source code available from https://github.com/kassonlab).

### Electron cryo-microscopy

Electron cryo-microscopy of endosome samples was performed at the University of Virginia Molecular Electron Microscopy core. Electron cryo-microscopy and analysis was performed as detailed in prior work [26]. Three microliters of sample were applied to a carbon-coated grid (2/2-4C C-flats; Electron Microscopy Sciences), blotted with filter paper, and plunge-frozen in liquid ethane. Samples were imaged in a Tecnai F20 Twin transmission electron microscope (FEI, Hillsboro, OR) at −180 C with a magnification of 29,000x, operating at 120 kV.

## Results and Discussion

We have established the following system to measure the kinetics of fusion between influenza virus and its physiological target membrane in a precise, chemically controllable manner. We fluorescently labeled influenza virus, infected cells with virus while preventing fusion, and isolated mannose 6-phosphate-receptor-positive (M6PR+) endosomes via gradient density separation. These endosomes were then immobilized in a microfluidic flow cell, and fusion was triggered by dropping the endosomal pH using an ionophore and buffer exchange. Lipid mixing kinetics between viral and endosomal membranes were measured as a function of time after pH drop. This system permits precise control of pH as well as characterization and manipulation of endosomal compartments not possible within living cells.

### Isolation of endosomes containing influenza virus

Baby hamster kidney (BHK) 21 cells were treated with 30 mM ammonium chloride prior to infection with X-31 (A/Aichi/68; H3N2) influenza virus at MOI of 1, to block endosomal acidification and subsequent viral fusion [27]. Cells were mechanically lysed, and M6PR+ endosomes were isolated by sucrose density gradient separation from the post nuclear supernatant of the infected cells. A cryo-EM image of an isolated endosome enclosing what is likely a virus is rendered in Figure 1b. Endosome fractions positive for both hemagglutinin and M6PR on immunoblot (Fig. S1) were biotinylated and surface-immobilized in a microfluidic flow cell passivated with PLL-PEG (Fig. 1c). This immobilization procedure yielded M6PR+ endosomes colocalizing with internalized Texas-Red-labeled virions upon immunostaining (Fig. 1d).

### Fusion between single viruses and endosomes reported by lipid mixing

In a variation on our previously reported procedure [18], fusion was triggered by buffer exchange to pH 5.0 inside the flow cell with the addition of the ionophore FCCP and simultaneous washout of ammonium chloride. This combination was designed to achieve rapid pH drop inside the endosomes [28,29] and was verified using pH-dependent fluorescence in synthetic liposomes (Fig. S2). Lipid mixing between individual encapsulated virions and the surrounding endosomal membranes was measured by fluorescence dequenching (see Fig. 2 for a sample trace), and individual waiting times between pH drop and fluorescence dequenching compiled into a cumulative distribution function (CDF) (Fig. 3). These measurements of isolated endosomes only assess lipid mixing directly, but based on similar ammonium chloride washout in cells and subsequent measurement of viral protein expression, full fusion likely occurs. Furthermore, fusion to any endosomal membrane within a single- or multi-vesicular body (see Fig. S3) will be sufficient to result in dequenching [30].

**Figure 2.**
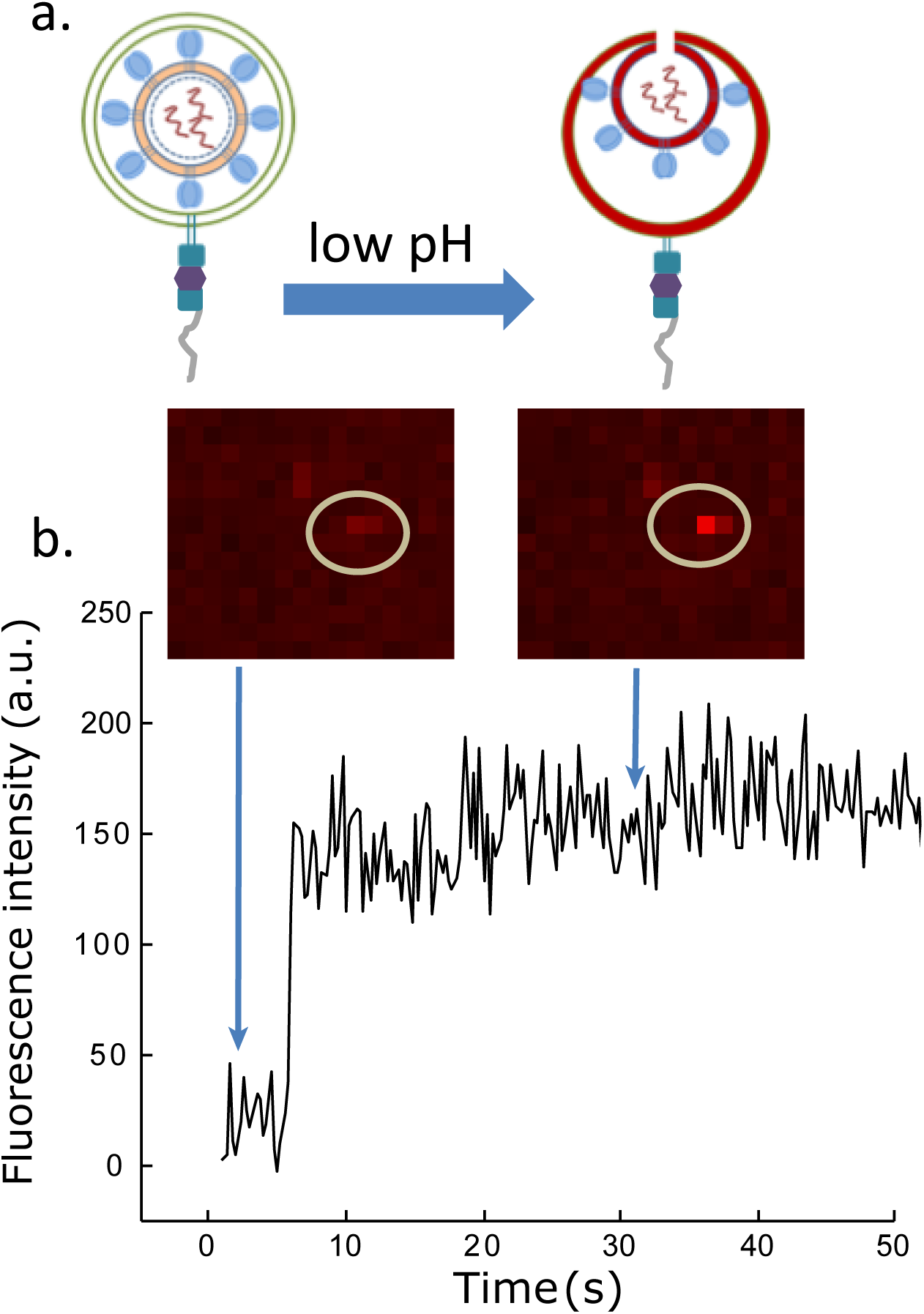
Hemi-fusion between viral and endosomal membranes. (a) Schematic representation of lipid mixing from viral to endosomal membranes at low pH. Lipid mixing is identified as the rapid increase in fluorescence due to dilutional dequenching of the lipid dye. This schematic is intended to illustrate the experiment rather than as a mechanistically specific model for viral fusion. (b) Representative intensity time trace of a single virus fusing with an endosome. Fluorescent images of the virus:endosome conjugate before and after pH change are shown. The waiting time is the time between pH drop and lipid mixing.

**Figure 3.**
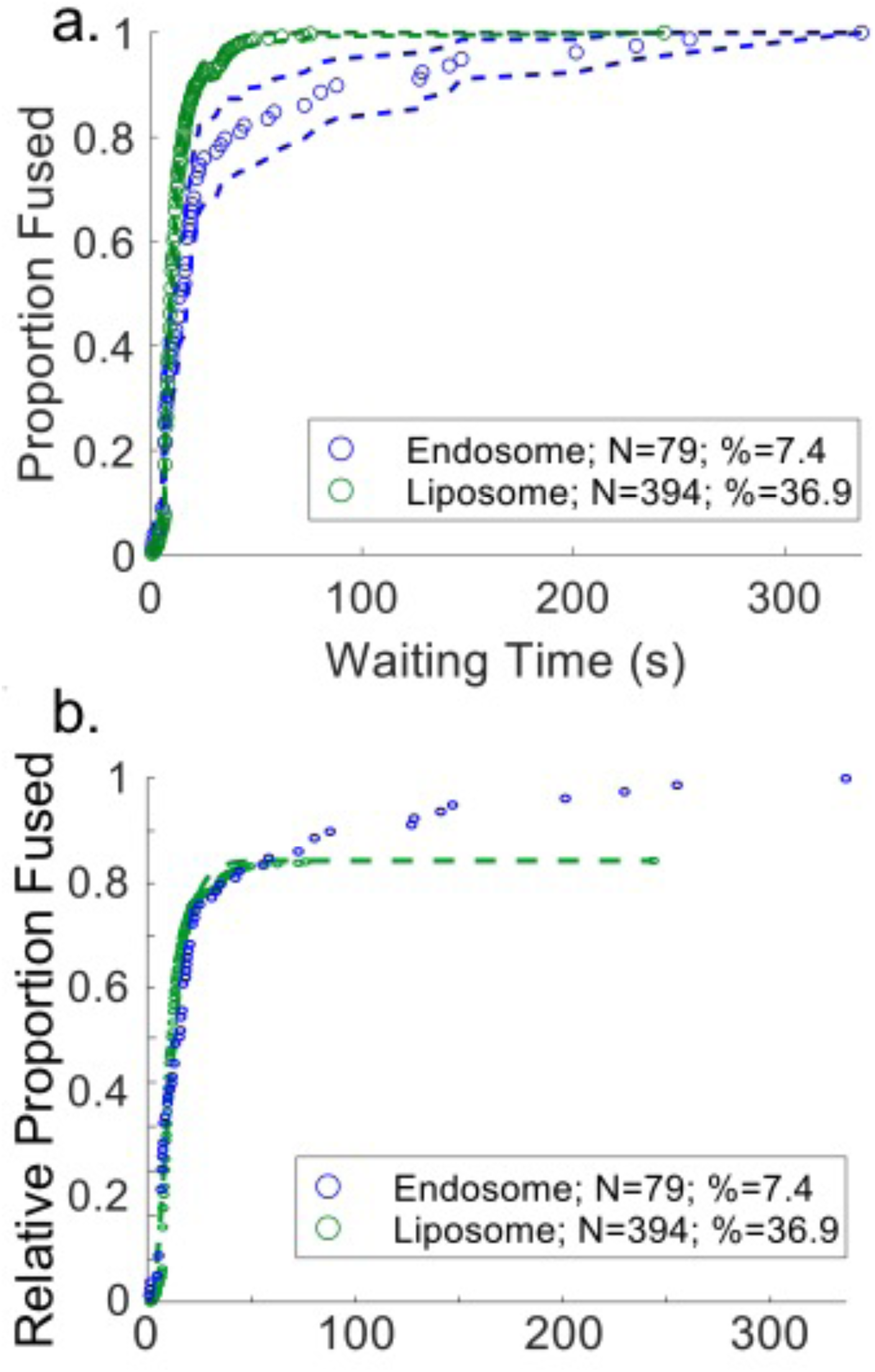
Comparison between lipid mixing kinetics of X-31 influenza virions bound to target liposomes by GD1a receptor and that of X-31 influenza virions with endosomes. Panel (a) shows cumulative distribution functions (CDF) of lipid-mixing waiting time distributions for single-virus:liposome (green) and single-virus:endosome (blue) fusion events, together with bootstrap resampling error estimates (95% confidence intervals). Panel (b) shows the same two distributions scaled to visualize the near-identical fast phase in the single-virus:endosome lipid mixing kinetics. Data represent the aggregate of two independent endosome preparations.

#### Kinetic similarity between virus-endosome and virus-liposome fusion

Model systems previously developed to study single-virus fusion kinetics have used planar bilayers or liposomes, typically of defined composition [13,16–19,31]. Our single-virus:endosome fusion platform enables assessment of whether those model approximations affect the rate-limiting steps for lipid mixing. We therefore compared the lipid-mixing kinetics of individual viruses fusing to the interior of endosomes against the kinetics of individual viruses fusing to the exterior of synthetic liposomes. Notwithstanding the curvature and compositional differences between endosomes and liposomes, the primary phase of lipid-mixing between influenza virus and endosomes overlaps well with virus-liposome fusion (Fig. 3). Fusion to endosomes was grossly biphasic, exhibiting a slower phase that do not overlap with the kinetics of fusion to liposomes. The overall agreement of the CDFs is consistent with recent work suggesting that lipid mixing rates are insensitive to membrane curvature [32]. Despite variation in size and curvatures among endosomes (Fig. S3), all present a negatively curved surface to viral fusion rather than the positively curved surface presented by the outside of liposomes.

Our findings support the hypothesis that influenza lipid-mixing kinetics are generally insensitive to target membrane curvature and that synthetic target membranes provide a good model for capturing the first-order fusion behavior of such viruses. The slower phase of lipid-mixing kinetics detected, however, does suggest the possibility of a sub-population of endosomes that may behave differently. In addition, the agreement in lipid-mixing kinetics does not exclude the possibility of a larger difference in kinetics in fusion pore opening and viral contents release between model target membranes and endosomes.

## Conclusions

Because viral fusion inside the cell is highly dependent on factors such as endosomal maturation kinetics and intracellular trafficking, chemically controllable platforms to measure entry of individual viruses are essential. Here, we have described the creation of such a platform to measure fusion kinetics of individual viruses within purified endosomes, preserving the physiological compartment for fusion but allowing precise time sequencing and permitting further chemical manipulation. We demonstrate that the kinetics of lipid mixing between influenza viruses and these endosomes recapitulates viral fusion to the exterior of model liposomes, with the addition of a second, slow-fusing population of virus:endosome conjugates. The endocytic fusion system recapitulates physiological curvature, composition, and asymmetry (modulo the perturbation provided by fusion arrest) and offers a platform to examine cellular restriction factors as well as pharmacological perturbation of cell metabolism and membrane composition.

## Acknowledgements

The authors acknowledge R. Dunning for initial pilot experiments, A. Sengar, A. Villamil Giraldo, J. White, and D. Castle for helpful advice and discussions, and J. White and D. Castle for the kind gift of reagents. Electron cryo-microscopy was performed by K. Dryden at the Molecular Electron Microscopy Core at the University of Virginia. This work was supported by a Wallenberg Academy Fellowship and Swedish Research Council grant 2017-04236 to P.M.K.

## Notes

### Competing Interest Statement

The authors have declared no competing interest.

