## Supplementary Figures for "Precise triggering and chemical control of single-virus fusion within endosomes"

### Measuring single-virus fusion within endosomes in a chemically controllable manner

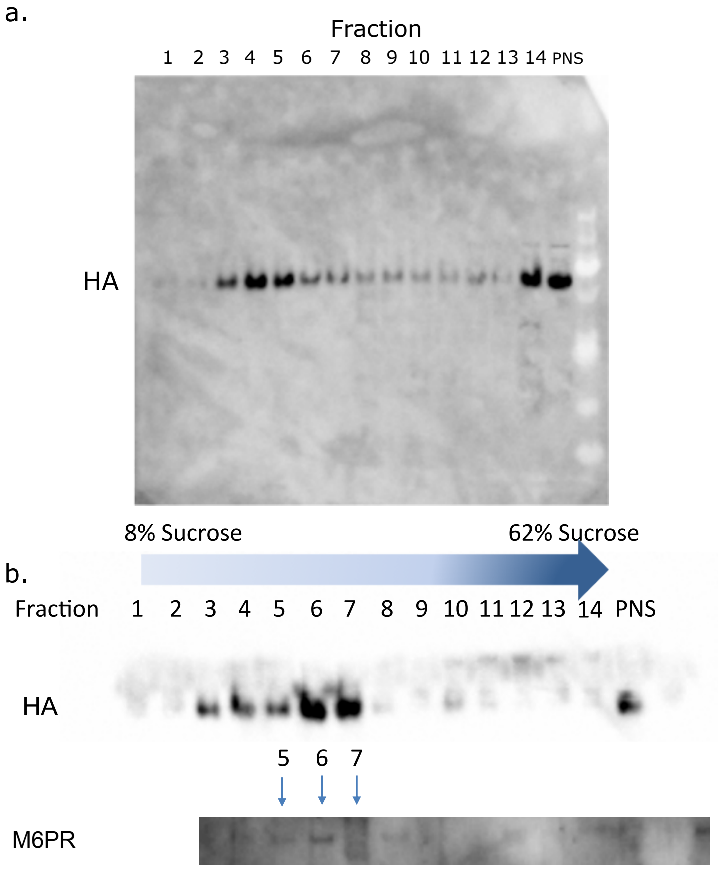


**Figure S1. Immunoblots of endosome fractions**. Rendered in panel (a) is an anti-hemagglutinin (HA) immunoblot of consecutive fractions from discontinuous sucrose gradient centrifugation. Rendered in panel (b) are two immunoblots from a sucrose gradient of a separate endosomal preparation with anti-hemagglutinin (HA) and anti-mannose-6-phosphate-receptor (M6PR) probed separately.

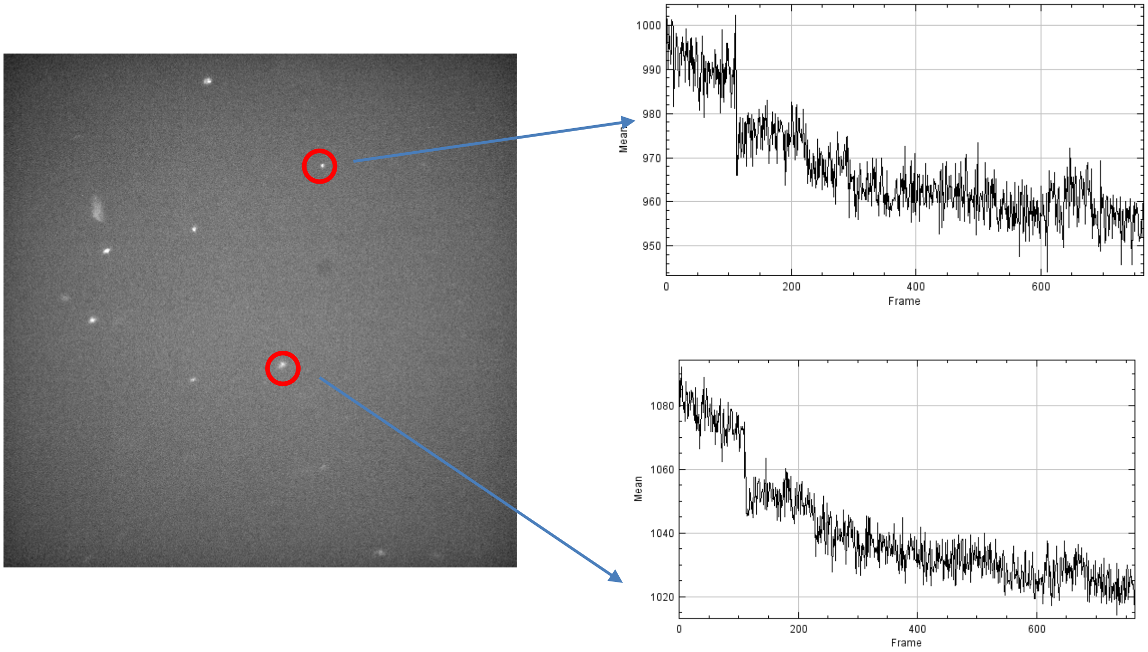


**Figure S2. Rapid luminal pH drop achieved by ionophores**. The left panel shows a fluorescence micrograph of vesicles labeled with carboxyfluorescein, a pH-sensitive dye, in their interior. On the right are two time traces from the indicated spots. Inflow of low-pH buffer (pH 5.0) with FCCP ionophore causes a rapid and synchronized drop in pH, as seen by the fluorescence drop superimposed on an overall bleaching profile. Carboxyfluorescein shows more bleaching under the imaging conditions used here than the Texas Red dye used for viral labeling.

a.

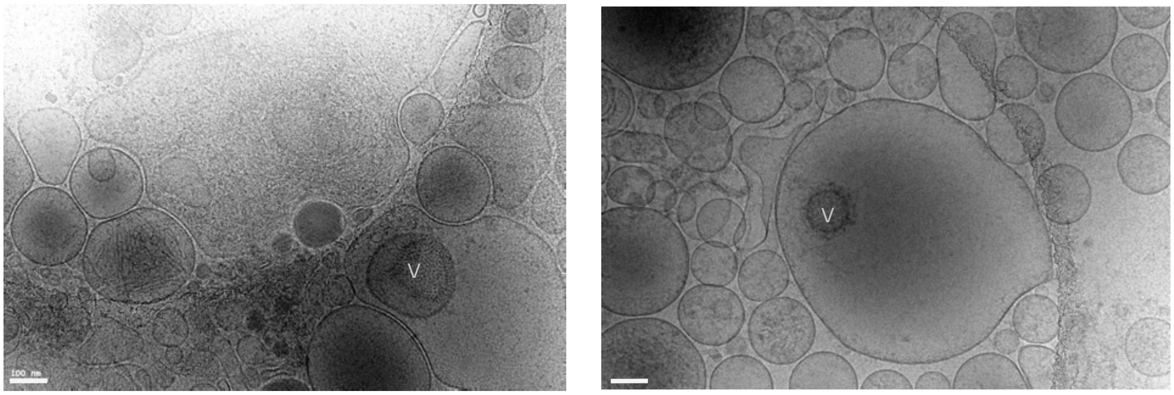


b.

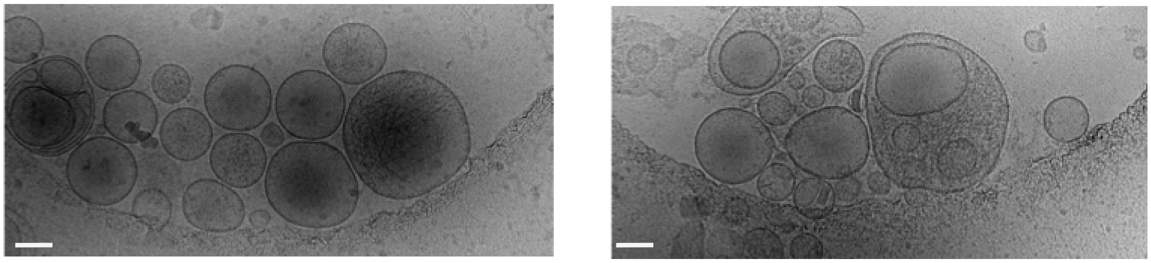

**Figure S3. Electron cryo-micrographs of endosomes with and without virus**. Electron cryo-micrographs are displayed in panel (a) with two enclosed viral particles, denoted by v. Cryo-micrographs of endosomes not containing viral particles are displayed in panel (b). Scale bars represent 100 nm.
